# Afucosylated maternal anti-dengue IgGs are a biomarker for susceptibility to dengue disease in their infants

**DOI:** 10.1101/565259

**Authors:** Natalie K. Thulin, R. Camille Brewer, Robert Sherwood, Stylianos Bournazos, Karlie G. Edwards, Nitya S. Ramadoss, Jeffery K. Taubenberger, Matthew Memoli, Prasanna Jagannathan, Sheng Zhang, Daniel H. Libraty, Taia T. Wang

**Affiliations:** Department of Medicine, Stanford University School of Medicine, Stanford, CA 94305, USA.; Proteomics Facility, Institute of Biotechnology, Cornell University, Ithaca, NY, 14853, USA.; The Laboratory of Molecular Genetics and Immunology, The Rockefeller University, 1230 York Avenue, New York, NY 10065, USA.; Department of Immunology and Rheumatology, Stanford University, Stanford, CA 94305, USA.; Viral Pathogenesis and Evolution Section, Laboratory of Infectious Diseases, Division of Intramural Research, National Institute of Allergy and Infectious Diseases, National Institutes of Health, Bethesda, Maryland, USA.; Department of Microbiology and Immunology, Stanford University, Stanford, California 94305, USA.; University of Massachusetts Medical School, Worcester, MA.; Chan Zuckerberg Biohub, San Francisco, CA 94518.

## Abstract

Infant mortality from dengue disease is a devastating global health burden that could be minimized with the ability to identify susceptibility for severe disease prior to infection. While most primary infant dengue infections are asymptomatic, maternally derived anti-dengue IgGs present during infection can trigger progression to severe disease through antibody-dependent enhancement mechanisms. Importantly, specific characteristics of maternal IgGs that herald progression to severe infant dengue are unknown. Here, we define ≥10% afucosylation of maternal anti-dengue IgGs as a biomarker for susceptibility of infants to symptomatic dengue infections. Mechanistic experiments show that anti-dengue afucosylation, a modification that enhances Fc affinity for the activating receptor FcγRIIIa, promotes infection of FcγRIIIa+ monocytes. FcγRIIIa signaling, in turn, enhances a post-entry step of dengue virus replication. These studies identify a biomarker that can be applied to reduce mortality associated with dengue viruses and define a mechanism by which afucosylated antibodies and FcγRIIIa enhance dengue infections.

## Introduction

Antibody-mediated inflammatory responses are critical in immunity against infectious organisms. These responses can promote pathogen clearance but can also exacerbate symptoms during infections (Bournazos et al., 2017). Antibody-mediated inflammation is triggered when pathogens or infected cells are bound by immunoglobulin G (IgG) antibodies, forming immune complexes that signal through Fc gamma receptors (FcγRs) on effector cells. The outcome of effector cell responses depends on the balance of activating to inhibitory (A/I) FcγR signaling that arises from interactions with Fc domains within immune complexes. One factor that modulates the ratio of A/I FcγR signaling is the glycosylation state of the IgG Fc domains within immune complexes. For example, sialylation of the Fc promotes anti-inflammatory effector responses while afucosylation of the Fc is pro-inflammatory due to increasing affinity of the Fc for the activating FcγRIIIa, found on NK cells, as well as on subsets of macrophages and monocytes (Anthony et al., 2011; Bournazos et al., 2017; Rafiq et al., 2013). The second major determinant of A/I FcγR signaling by immune complexes is the distribution of IgG subclasses within the complex with IgG1 being the dominant subclass promoting pro-inflammatory responses and IgG2 signaling through the inhibitory FcγR (Pincetic et al., 2014).

Dengue virus infections are unusual in that non-neutralizing anti-dengue virus IgGs can play a central role in triggering progression to the severe forms of disease through antibody-dependent enhancement (ADE) mechanisms (Anderson et al., 2014; Burke et al., 1988; Chau et al., 2009; Guzman et al., 1990; Halstead et al., 1970; Katzelnick et al., 2017; Libraty et al., 2009; Sangkawibha et al., 1984; Wang et al., 2017). ADE can occur in the presence of reactive, non-neutralizing IgGs, as are found in secondary heterologous dengue infections or in primary infections in infants of dengue-immune mothers due to acquisition of anti-dengue virus IgGs during gestation (Halstead et al., 1970; Kliks et al., 1988; Simmons et al., 2007). These antibodies are thought to promote disease by forming immune complexes with the virus that modulate infection in FcγR-bearing cells, primarily monocytes and macrophages, leading to higher viral titers and altered cytokine production during infection (Aye et al., 2014; Durbin et al., 2008; Kou et al., 2008; Thein et al., 1997). Still, the vast majority of dengue infections that occur in the presence of non-neutralizing IgGs are asymptomatic and specific features of antibodies that enhance one’s basal susceptibility for dengue disease are unknown.

Importantly, mortality rates in severe dengue can exceed 20% when patients are not hospitalized, but can be reduced to <1% with in-patient care (Anderson et al., 2014; Gordon et al., 2013). Therefore, the identification of biomarkers for increased baseline susceptibility to dengue disease could dramatically reduce mortality associated with these viruses by enabling early hospitalization of those at highest risk for disease progression. Here, we have conducted a global analysis of anti-dengue IgGs from mothers of infants with known disease severity during primary dengue infections in order to identify features of maternal IgGs that portend dengue disease risk in their infants. Primary dengue infection in infants is a model system to study the role of pre-infection IgGs on dengue disease susceptibility because maternal IgGs, but no other antibody isotypes, are transferred to the fetus during gestation. Further, in primary infant dengue infections all anti-dengue IgGs present during infection are maternally-derived as the infant has not been previously infected. In contrast to studying secondary heterologous infections, primary infection in infants is also not confounded by the potential of contributions from other adaptive immune elements (T cells, etc.) that are present in dengue-experienced individuals (Nicoara et al., 1999; Techasena et al., 2007). Thus, the maternal anti-dengue repertoire can be studied as a surrogate for the pre-infection IgG repertoire in their infants.

Here we show that IgGs modified by Fc glycoforms that modulate inflammatory effector functions, sialylation and fucosylation, are stable in abundance over time and are transferred neutrally across the placenta from maternal to cord blood. Further, elevated levels of afucosylated maternal anti-dengue IgGs predicts susceptibility of their infants to disease during primary dengue infections. Because afucosylation enhances affinity of the Fc for FcγRIIIa, this finding suggests a role for FcγRIIIa in the pathogenesis of dengue disease. We therefore performed studies to dissect the mechanisms by which FcγRIIIa and afucosylated IgGs impact dengue virus infections. Our findings identify a novel FcγRIIIa-mediated ADE pathway by which dengue virus infections can be enhanced. Thus, we elucidate a mechanistic basis for differential susceptibility of infants to dengue disease and identify a specific biomarker that can be applied to improve the clinical management of dengue patients.

## Results

### IgG glycoforms are stable over time and neutrally transferred across the placenta

For IgG Fc glycans to be a useful biomarker for disease susceptibility, their abundance would have to be stable over time and withstand immune perturbation. To evaluate this, we performed a longitudinal study of anti-hemagglutinin (HA) IgGs from healthy adults in a North American cohort who were challenged with influenza A virus (Memoli et al., 2015). Due to regular exposures to influenza viruses in this region, anti-HA IgGs are nearly always present in adults and are thus a practical specificity for this analysis. To define durability of Fc glycoforms in humans, we performed an analysis of longitudinal samples over a period of 60 days after influenza A challenge in this cohort. The abundance of sialylated and afucosylated glycans was remarkably durable over this period despite influenza virus challenge (Figure 1A).

**Figure 1.**
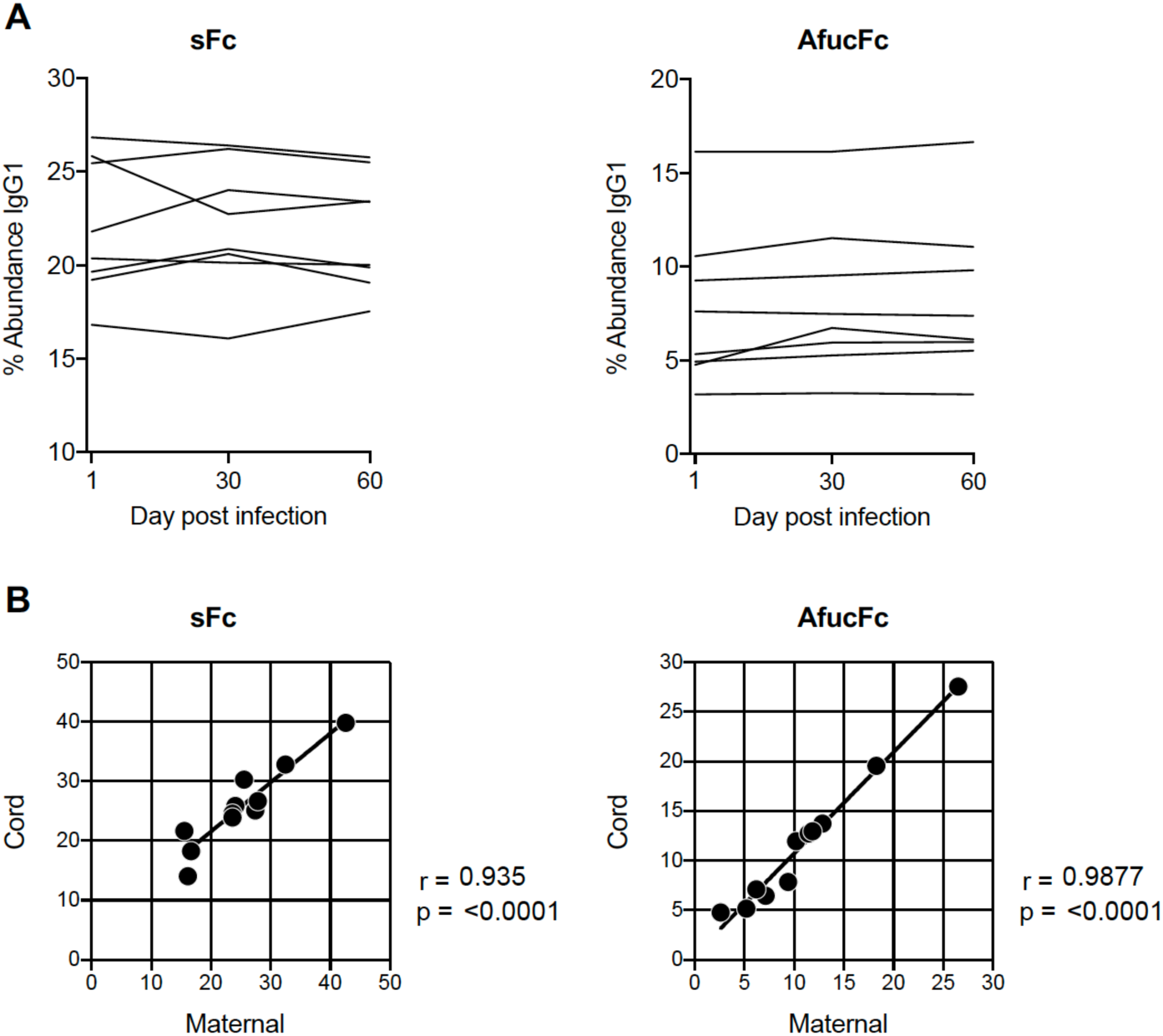
Stability of Fc glycans over time and neutral transfer across the placenta. **A)** The abundance of sialylated (sFc) and afucosylated (afucFc) glycans is remarkably stable over 60 days in healthy adults challenged with influenza A virus Influenza virus challenge took place on day 1. Longitudinal samples from days 1, 30 and 60 post challenge are shown. **B)** The abundance of sialylated and afucosylated Fc glycans is highly correlated between maternal and cord blood lgGs demonstrating neutral transpiacental transfer.

We next asked whether maternal IgGs can be used as a surrogate for neonatal IgGs with respect to Fc glycosylation. For this to be true, IgGs would have to be transferred neutrally across the placenta despite different Fc glycosylation. To address this, we studied the abundance of sialylated and afucosylated IgGs in paired maternal and cord bloods from a Ugandan cohort (Jagannathan et al., 2018). The abundance of maternal IgGs with sialylated or afucosylated Fc glycoforms was highly predictive of their abundance in matched cord blood (p<0.0001) (Figure 1B). Overall, these data show that the abundance of IgG Fc glycoforms is durably heterogeneous among people despite immune perturbation and the maternal repertoire of sialylated and afucosylated IgG Fc glycans determines the neonatal repertoire.

### Elevated afucosylation of maternal anti-dengue IgGs predicts susceptibility to infant dengue disease

To define specific maternal anti-dengue IgG features that herald risk for infant dengue disease, we performed a global analysis of anti-dengue envelope (E) IgGs in mothers of infants with symptomatic or asymptomatic primary dengue infections. To do this, we studied samples from a prospective cohort in San Pablo, Laguna, Philippines, a dengue-endemic region. After delivery, infants were followed rigorously for primary dengue infections including documentation of subclinical infections evidenced by seroconversion (Capeding et al., 2010; Libraty et al., 2009).

Given the role of FcγRs in ADE of dengue virus infections, we first asked whether any maternal anti-E Fc glycosylation patterns could predict susceptibility to infant dengue disease. Mothers of infants with clinically significant (symptomatic) primary dengue virus infections had significantly elevated levels of afucosylated anti-E IgG compared with mothers of infants who had subclinical primary dengue virus infections (Figure 2A). Maternal, afucosylated anti-dengue Fc glycans ≥10% in abundance was a significant predictor of symptomatic infant primary dengue infections (positive predictive value: 87.5% (95% CI 52.9%-99.3%)). No significant differences in other Fc glycoforms were found, including bisecting N-acetylglucosamine (GlcNAc), a modification that may also impact Fc-FcγRIIIa interactions, though to a lesser extent than afucosylation (Figure 2B)(Shinkawa et al., 2003).

**Figure 2.**
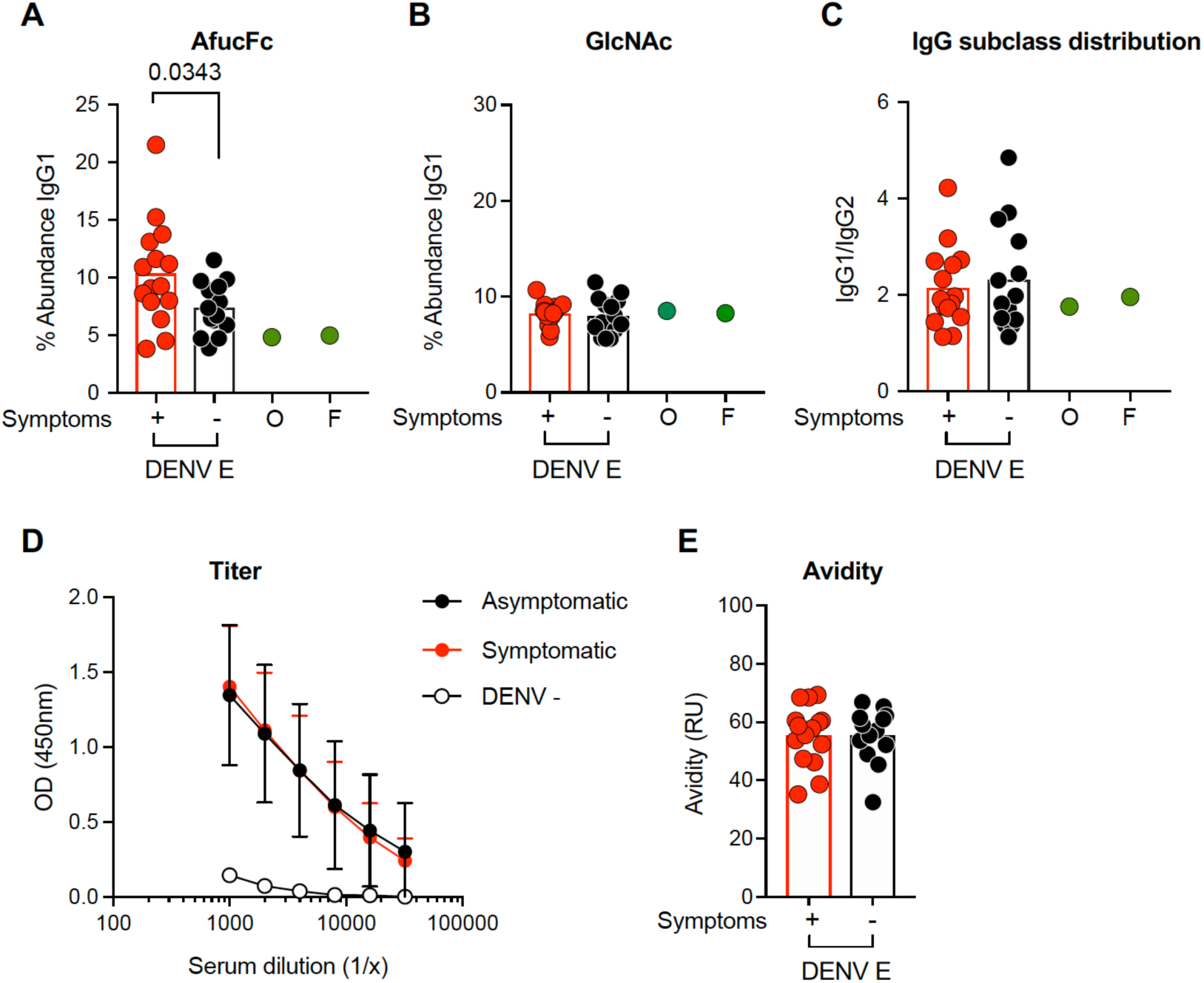
Elevations in maternal, afucosylated anti-DENV lgGs predict symptomatic, primary infant dengue virus infections. **A)** ≥10% afucosylation of maternal anti-dengue E (DENV E) IgG1 predicts symptomatic infection in their infants (positive predictive value: 87.5% (95% CI 52.9%-99.3%)). Symptomatic ^(+)^; asymptoniatic (-). Two commercial IVIg preparations are shown (O and F) **B)** Other Fc glycoforms were not significantly different between mothers of infants with symptomatic and asymptomatic dengue infections, induding bisecting GIcNAc. a modification that has been reported to impact FcγRIIIa interactions. There were no differences the maternal anti-E lgG **C)** subclass distribution, **D)** titer on dengue-infected cells or **E)** avidity between mothers of infants who developed symptomatic or asymptomatic primary dengue infections.

To determine whether other anti-E features, aside from Fc afucosylation, herald symptomatic infant dengue infections, we studied the other antibody characteristics that can impact the A/I ratio of Fc-FcγR interactions. Symptomatic or asymptomatic primary infant dengue infections could not be predicted by the distribution of activating to inhibitory maternal anti-E subclasses (IgG1/IgG2) (Figure 2C). In addition, the titer of serum anti-E IgGs was not different between mothers of infants with symptomatic or asymptomatic dengue infections (Figure 2D). To determine whether afucosylation could be a surrogate for antibody titer, we compared the anti-dengue binding titer in subjects with ≥10% or <10% Fc afucosylation and observed no difference between groups (Figure S1A). Finally, there was no difference in the binding avidity of anti-E IgGs between mothers of infants with symptomatic or asymptomatic primary dengue infections (Figure 2E).

Because anti-NS1 IgGs can modulate the severity of dengue infections we further assessed whether the maternal titer of anti-NS1 IgGs predicted symptomatic dengue infections in their infants and found no correlation (Figure S1B). In addition, there were no differences between anti-NS1 Fc glycoforms from mothers of infants with symptomatic or asymptomatic primary dengue infections (Figure S1C). There were also no differences in Fc glycoforms of total IgGs between mothers of infants with symptomatic or asymptomatic primary dengue infections demonstrating the specificity of anti-E afucosylation as a predictor of symptomatic infant dengue infections (Figure S1C).

Overall, we find that ≥10% afucosylated glycans on maternal anti-E IgGs is a biomarker for susceptibility of their infants to clinically significant primary dengue infections. No other antibody correlates of symptomatic infant dengue infections were observed. Clinical utility of this biomarker is supported by the finding that afucosylated IgGs are durable in abundance (Figure 1) and may therefore be measured within a window of time after delivery to assess susceptibility of their newborns to dengue disease.

### Afucosylated anti-dengue virus IgGs skew infection of monocytes towards those expressing FcγRIIIa

Afucosylation of the Fc increases its affinity for the activating FcγR, FcγRIIIa. This receptor, while present on macrophage and pro-inflammatory monocyte subsets that are associated with dengue disease, is not expressed on human monocytic or other granulocytic lineage cell lines often used to study ADE of dengue virus infections such as THP-1, K562 or U937 cells (Fleit et al., 1982; Jones et al., 1985; Tridandapani et al., 2002; van de Winkel and Anderson, 1991). These cells lines express the high-affinity activating FcγR, FcγRI (except for K562 cells) which is occupied by monomeric IgG at baseline; additionally, they express the low-affinity activating FcγR, FcγRIIa as well as the low-affinity inhibitory FcγR, FcγRIIb. To study how anti-E IgG Fc afucosylation and FcγRIIIa impact ADE, we generated human U937 monocytes expressing different combinations of the activating FcγRs, FcγRIIa and FcγRIIIa (Figure S2). While FcγRIIa has been previously identified as a key receptor in mediating aspects of ADE (Boonnak et al., 2013; Chawla et al., 2013; Rodrigo et al., 2006), the role of FcγRIIIa in ADE of dengue virus infection has not been defined.

The role of afucosylation of anti-E IgGs in ADE of dengue infection was first studied by infecting cells with dengue immune complexes generated from fucosylated or afucosylated anti-dengue E mAb D23-5G2D2 (mAb 235) (Sasaki et al., 2013) (Figure S3). Cells expressing FcγRIIa^+^FcγRIIIa^-^ or FcγRIIa^+^FcγRIIIa^+^, which are physiologically relevant expression patterns for these receptors, were tested. No difference was observed in the percent infection of pure populations of these cells as a function of the anti-E fucosylation state. However when infections were performed in a mixed monocyte population representing the approximate distribution of FcγRIIa^+^FcγRIIIa^+^ and FcγRIIa^+^FcγRIIIa^-^ subsets *in vivo*, afucosylation induced preferential infection of FcγRIIa^+^FcγRIIIa^+^ monocytes (Figure 3A,B, Figure S4) (Boyette et al., 2017). To determine whether the fine specificity of anti-E IgGs might impact this finding, we expressed additional anti-E mAbs 747(4) B7 (B7), 2D22, and 753(3) C10 (C10) as human IgG1s with fucosylated or afucosylated Fc glycans (Figure S3, Table S1) (Rouvinski et al., 2015; Swanstrom et al., 2016). As with mAb 235, we found that significantly more FcγRIIa^+^FcγRIIIa^+^ monocytes were infected when immune complexes were generated from afucosylated anti-E mAbs (Figure 3B), demonstrating that the fucosylation status of anti-E antibodies can shift the distribution of infected monocytes toward those expressing FcγRIIIa.

**Figure 3.**
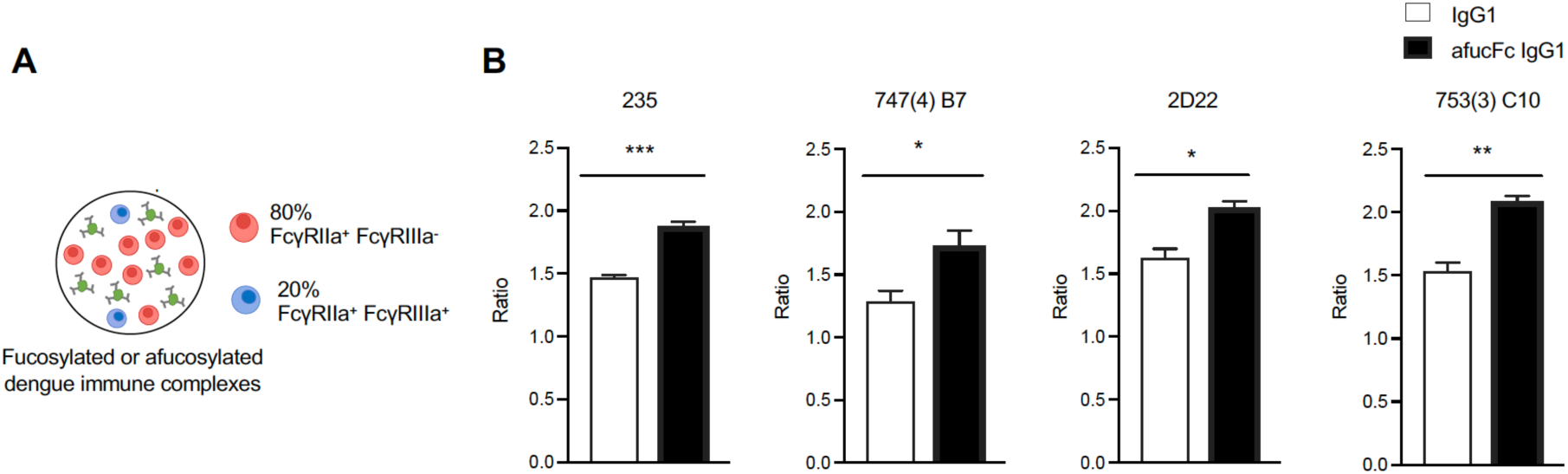
Afucosylation of anti-dengue IgGs increases infection in FcγRIIIa^+^ monocytes. **A)** FcγRIIa^+^FcγRIIIa^+^ and FcγRIIa^+^FcγRIIIa^-^ cells were mixed at a ratio of 20:80, the approximate physiologic ratio of these monocyte subsets *in vivo*. The mixed cell populations were infected with fucosylated or afucosylated dengue inmune complexes and the ratio of infection in FcγRIIa ^+^FcγRIIIa^+^:FcγRIIa^-^FcγRIIIa^+^ was determined after 24 hours. **B)** Infection with afucosylated immune complexes generated with different anti-E mAbs 235, B7, 2D22 and C10) skewed infection towards FcγRIIa^+^FcγRIIIa^+^ monocytes.

### FcγRIIa and FcγRIIIa have distinct roles in ADE of dengue virus infection

The findings that elevated afucosylated glycans on anti-E IgG1 predicts risk for symptomatic dengue infection in infants and can bias infection towards FcγRIIIa-expressing cells suggests that FcγRIIIa plays a role in the pathogenesis of dengue disease. To study the respective roles of FcγRIIa and FcγRIIIa in ADE of dengue virus infections, we infected U937 monocytes expressing different combinations of these receptors with dengue virus/mAb 235 immune complexes. When we compared infection in cell lines expressing FcγRIIa^-^FcγRIIIa^-^, FcγRIIa^-^FcγRIIIa^+^, FcγRIIa^+^FcγRIIIa^-^, or FcγRIIa^+^FcγRIIIa^+^ we found that expression of FcγRIIIa alone had no significant impact on infection (FcγRIIa^-^FcγRIIIa^-^ vs. FcγRIIa^-^FcγRIIIa^+^ cells). Interestingly however, expression of FcγRIIIa in the presence of FcγRIIa amplified infection by over 70% as compared to infection of monocytes expressing only FcγRIIa (Figure 4A). That FcγRIIIa only contributed to ADE in the presence of FcγRIIa demonstrated distinct roles for these receptors in ADE of infection by dengue virus immune complexes.

**Figure 4.**
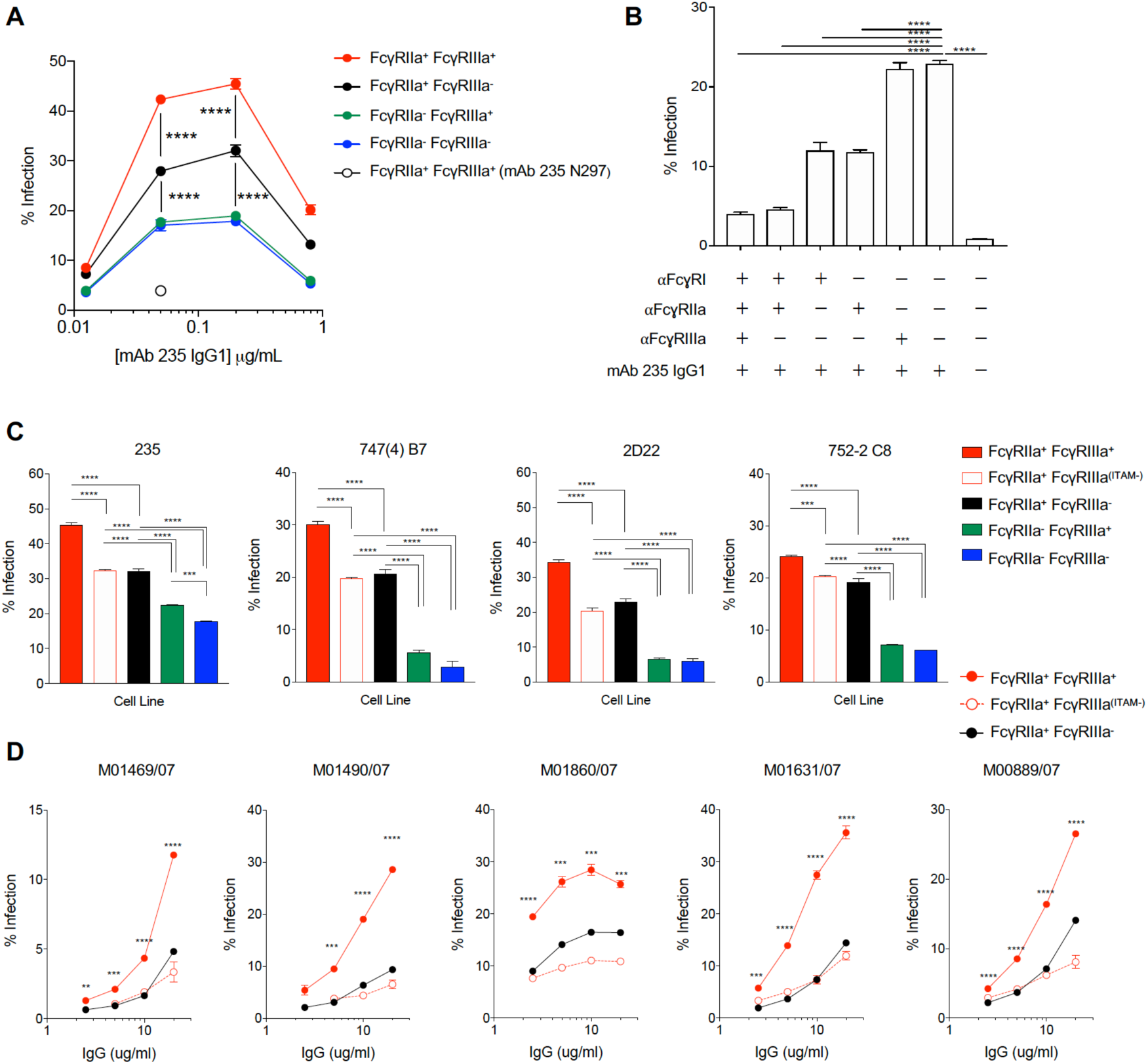
FcyRl and FcγRIIa mediate enhanced entry of DENV immune complexes while FcγRIIIa signaling mediates a post-entry step in ADE. **A)** Immune complexes composed of human IgG1 mAb 235 and DENV2 were used to infect the panel of U937 cell lines. Infectability of the FcγRIIa^-^FcγRIIIa^-^ (blue) and FcγRIIa^-^FcyRIII^+^ (green) cell lines was similar, demonstrating that expression of FcγRIIIa alone did not impact infection In contrast, expression of FcγRIIa alone (FcγRIIa^+^FcγRIIIa^-^, black) increased infectability and co-expression of FcγRIIa and FcγRIIIa (FcγRIIa^+^FcγRIIa^+^, red) further enhanced infection. Infection in the presence of mAb 235 N297A, an Fc mutation that abrogates FcyR binding, is shown in the unfilled circle. **B)** To determine whether the presence of FcγRIIIa enhances infectability of FcγRIIa^+^ FcγRIIIa^+^ cells by increasing virus binding to host cells or virus entry. FcyRs were individually blocked during infection. Blocking FcγRIIIa did not impact infection. **C)** Infection by mAb 235 dengue inmune complexes was reduced in the FcγRIIa^+^FcγRIIIa^(ITAM-)^ cells which lack FcγRIIIa signaling capacity. Infection by dengue immune complexes generated from anti-envelope mAbs 747(4) B7, 2D22 or 752-2 C8 followed the pattern of being increased in FcγRIIa^+^FcγRIIIa^-^cells over FcγRIIa-defldent cells, but was maximally enhanced by co-expression of FcγRIIa and FcγRIIIa. Enhancement in FcγRIIa^+^FcγRIIIa^+^ cells required FcγRIIIa signaling in all cases **D)** Infection by human polyclonal dengue immune complexes was maximal in FcγRIIa^+^FcγRIIIa^+^ monocytes and FcγRIIIa signaling was required for enhancement in all cases.

### FcγRIIa mediates entry in ADE while FcγRIIIa signaling mediates post-entry enhancement of dengue infection

To begin to define the roles of FcγRIIa and FcγRIIIa in ADE, we probed the contribution of these receptors in entry of dengue virus immune complexes. To do this, we used well described blocking mAbs against the activating FcγRs, FcγRI, FcγRIIa or FcγRIIIa during infection of FcγRIIa^+^FcγRIIIa^+^ monocytes (Mandelboim et al., 1999). The presence of blocking mAbs specific for either FcγRI or FcγRIIa during incubation of cells with virus immune complexes inhibited infection by approximately 50% whereas blocking FcγRIIIa alone did not inhibit infection (Figure 4B). Combining blocking mAbs specific for FcγRI and FcγRIIa inhibited infection by ∼80% and inclusion of the FcγRIIIa blocking mAb did not further diminish infection (Figure 4B). These results are consistent with prior studies implicating FcγRI and FcγRIIa in virus entry during ADE of dengue infections (Boonnak et al., 2013; Kou et al., 2008; Littaua et al., 1990; Rodrigo et al., 2006) and show a role for FcγRIIIa in enhancing dengue infection through a separate, post-binding and entry mechanism.

To determine whether receptor signaling was required for FcγRIIIa-mediated ADE, we assessed ADE in cells expressing signaling competent (FcγRIIa^+^FcγRIIIa^+^) or signaling null (FcγRIIa^+^FcγRIIIa^(ITAM-)^) versions of FcγRIIIa (Figure S2B). The FcγRIIa^+^FcγRIIIa^(ITAM-)^ cells express a variant of FcγRIIIa that does not associate with the Fc receptor gamma chain, and thus lacks ITAM signaling (Blazquez-Moreno et al., 2017). Loss of FcγRIIIa-mediated enhancement was observed in the FcγRIIIa signaling-null cell line (Figure 4C). To exclude potential Fab-specific effects, we extended our study by including additional anti-E mAbs (B7, 2D22, and 752-2 C8) (Table S1) (Dejnirattisai et al., 2015; Rouvinski et al., 2015; Swanstrom et al., 2016). As with mAb 235, maximal infection was observed in FcγRIIa^+^FcγRIIIa^+^ cells and this FcγRIIIa-mediated enhancement was dependent on FcγRIIIa signaling (Figure 4C). To better characterize the impact of FcγRIIIa signaling on infection, we next studied polyclonal IgGs from five sera of mothers from the San Pablo cohort who live in a dengue-endemic region (Libraty et al., 2009). Maximal enhancement of dengue virus infection by polyclonal IgGs required signaling by FcγRIIIa in all cases and single-cycle infection was enhanced up to 300% in FcγRIIa^+^FcγRIIIa^+^ monocytes compared with those expressing only FcγRIIa (Figure 4D). Overall, these studies demonstrated that FcγRIIIa signaling can contribute to post-entry enhancement of dengue infection in monocytes.

### Post-entry enhancement of dengue infection requires the calcineurin signaling network

Engagement and crosslinking of the activating Type I FcγRs, FcγRI, FcγRIIa and FcγRIIIa by immune complexes triggers phosphorylation of ITAM domains by Src family kinases and intracellular calcium flux (Bournazos and Ravetch, 2015). Calcium flux, in turn, activates calcineurin, a serine/threonine-protein phosphatase which recognizes a host of substrates, scaffolds and regulator proteins through specific docking motifs (Roy and Cyert, 2009; Roy et al., 2007).

To dissect how enhanced ITAM signaling through FcγRIIIa activity contributes to ADE of dengue infection, cells were treated with inhibitors of the ITAM signaling pathway prior to infection. First, FcγRIIa^+^FcγRIIIa^+^ monocytes were treated with PRT-060318 (PRT), an inhibitor of Syk tyrosine kinase activity which is required for initiation of signaling through ITAM domains (Lowell, 2011; Reilly et al., 2011). Treatment with PRT inhibited dengue infection (Figure 5A), supporting a requirement for ITAM signaling in ADE of infection.

**Figure 5.**
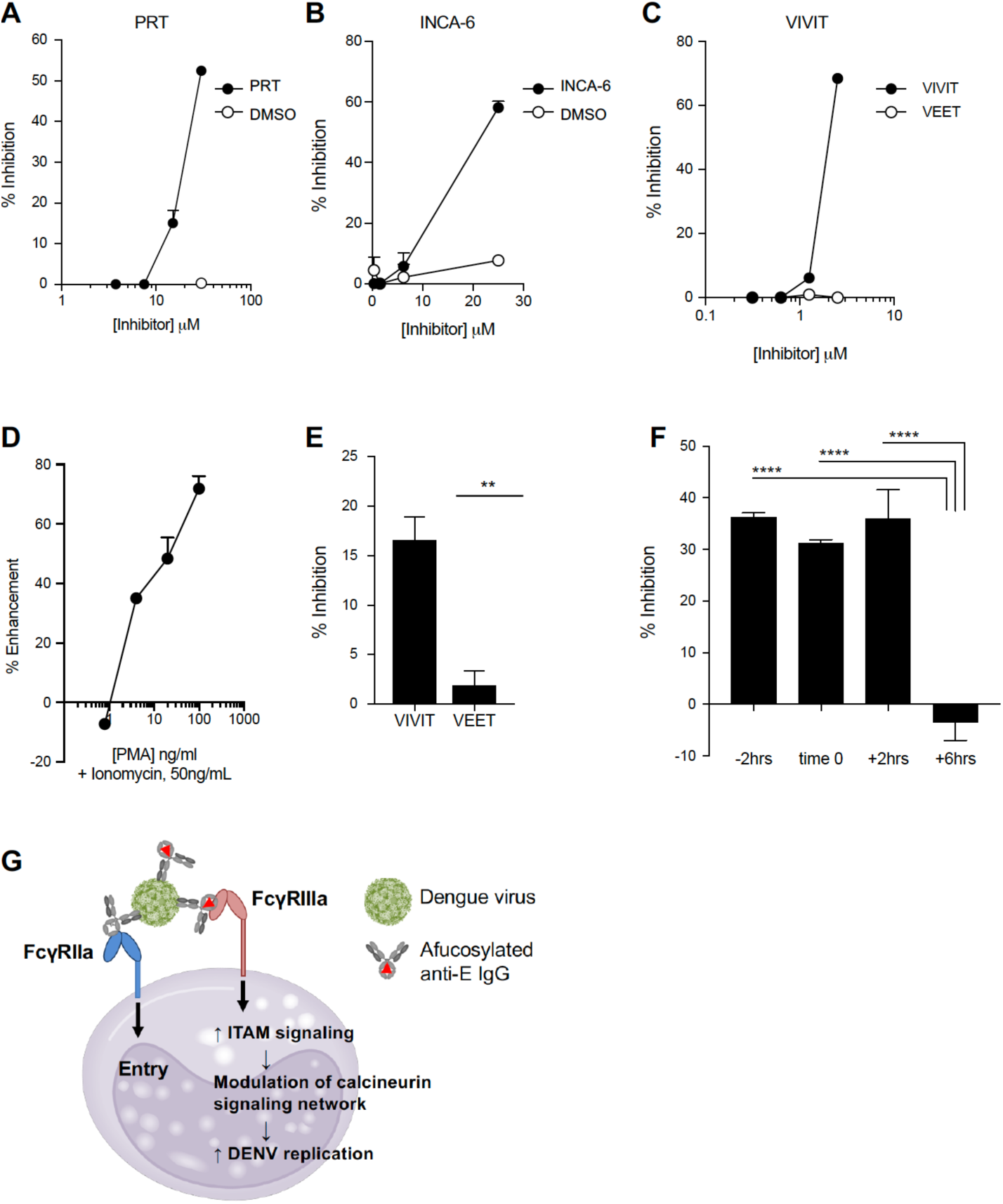
Inhibition of dengue infection through targeted inhibition of ITAM signaling pathway elements. **(A)** A small-molecule that targets Syk tyrosine kinase (Syk TK), PRT, inhibited dengue infection in FcγRIIa^+^FcγRIIIa^+^ monocytes. **(B)** A small molecule inhibitor, INCA-6, and **(C)** a peptide inhibitor, 11R-VIVIT (VIVIT), which block calcineurin-substrate interactions at the PxIxIT docking site inhibited dengue infection. A control peptide containing the amino acids found in VIVIT but in a scrambled sequence (VEET) did not inhibit infection. **(D)** Activation of NFAT-regulated genes by treatment of FcγRIIa^+^FcYRIIIa^-^ U937 cells with phorbol ester (PMA) plus ionomycin enhanced dengue infection. **(E)** Enhanced infection after PMA/ionomycin treatment could be inhibited by the inhibitor VIVIT but not by VEET. **(F)** Treatment of cells up to 2 hours following dengue infection with VIVIT prevented replication. **(G)** Model of role for FcγRIIa and FcγRIIIa in ADE of dengue infection in U937 monocytes. While FcγRIIa supports a majority of entry, FcγRIIIa signaling increases cellular ITAM signaling, in turn, modulating the calcineurin signaling network including NFAT-regulated genes which supports a post-entry step of dengue virus replication.

Next, to test the hypothesis that the calcineurin-substrate interactions may be required for ADE, we assessed the impact of inhibitors of this network on ADE of dengue infection. The small molecule inhibitor INCA-6 (Bretz et al., 2013) and the peptide inhibitor 11R-VIVIT (VIVIT) (Aramburu et al., 1999) act selectively to prevent the association of calcineurin with substrates utilizing a PxIxIT docking motif (Roy et al., 2007). These inhibitors prevented infection in a concentration dependent manner demonstrating a role for calcineurin network signaling in supporting dengue virus replication (Figure 5B,C). NFAT transcription factors are one calcineurin substrate that utilize the PxIxIT docking site and can be activated following FcγRIIIa cross-linking (Aramburu et al., 1995; Roy et al., 2007). NFAT transcription factors regulate an array of cellular genes (Crabtree and Olson, 2002; Feske et al., 2000; Rao et al., 1997). To test the hypothesis that activation of NFAT transcription factors during ADE supports dengue infection, we determined how direct activation of this pathway impacts infection of U937 cells with dengue immune complexes. Cells were treated with ionomycin and phorbol ester (PMA) which increase intracellular calcium levels and expression of an NFAT transcriptional co-regulator, activating protein 1, respectively (Hooijberg et al., 2000; Willingham et al., 2005). Pre-treatment of cells with PMA and ionomycin significantly enhanced infection by dengue immune complexes (Figure 5D). The enhanced infection could be inhibited by the VIVIT inhibitor which blocks the NFAT docking site on calcineurin, but not by the scrambled peptide control, VEET (Figure 5E).

To determine the stage of infection impacted by calcineurin network activity we performed time-course infections in which FcγRIIa^+^FcγRIIIa^+^ cells were treated with the VIVIT inhibitor 2 hours prior, during, 2 hours after, or 6 hours after infection with dengue virus immune complexes. Consistent with the post-entry enhancement arising from FcγRIIIa ITAM signaling, treatment of cells with the inhibitor up to 2 hours after infection prevented dengue replication (Figure 5F).

Together, these data support a model whereby maximal ADE of dengue virus infection is achieved through cooperativity between the low-affinity activating FcγRs, FcγRIIa and FcγRIIIa. While FcγRIIa acts in virus immune-complex entry, increased ITAM signaling through FcγRIIIa interactions supports post-entry viral replication through modulation of the calcineurin signaling network (Figure 5G).

## Discussion

Studies here show that the basal abundance of maternal anti-E afucosylation is a risk factor for susceptibility to disease upon dengue infection in their infants. To our knowledge, this constitutes the first demonstration of pre-infection IgG Fc glycosylation impacting susceptibility to an infectious disease. Further studies will determine whether baseline IgG glycosylation patterns may be useful in predicting risk for dengue disease after dengue vaccination, potentially enabling refinement of vaccination strategies (Anderson et al., 2018).

As mortality rates in severe dengue cases can be nearly eliminated with intensive clinical monitoring (Anderson et al., 2014; Gordon et al., 2013), the biomarker for susceptibility to infant dengue disease that is identified here can guide the development of tools to improve the clinical management of dengue patients. The potential for clinical utility of IgG Fc glycoforms as a predictive biomarker is bolstered by their stability in human serum over time and despite influenza virus challenge.

We have previously observed that disease severity during acute secondary dengue infections is impacted by elevations in afucosylated anti-dengue IgGs (Wang et al., 2017). Here, we demonstrate that this pro-inflammatory Fc glycoform can be utilized to predict susceptibility to dengue disease prior to infection. Together, these studies reveal a role for pro-inflammatory IgG Fc glycoforms in determining dengue disease susceptibility through modulating the progression of infection. Previous studies of acute human dengue infections have correlated increased infection in monocytes with increased disease severity and expansion of FcγRIIIa-expressing monocytes, specifically, with increased viral load (Kwissa et al., 2014; Srikiatkhachorn et al., 2012). The finding that the expansion of FcγRIIIa^+^ monocytes correlates with high viral load suggests that activation and/or infection of this specific cell subset may play a pivotal role in dengue pathogenesis (Kwissa et al., 2014).

In these experiments, we have used percent infection as the readout of activity conferred by anti-dengue Fc afucosylation and of FcγRIIIa in ADE of dengue infections. Fc afucosylation and FcγRIIIa signaling activity may impact dengue disease pathogenesis through multiple mechanisms including modulating virus production and activation and cytokine production by FcγRIIIa-expressing cells, including NK cells.

Overall, these studies support a model whereby basal heterogeneity in maternal IgG Fc fucosylation drives susceptibility to infant dengue disease severity through FcγRIIIa signaling-dependent pathways (Figure S5). Increased production of pro-inflammatory/afucosylated immune complexes increases the A/I signaling ratio during infection, modulating the calcineurin signaling network which supports dengue infection after virus entry. Specifically, calcineurin interactions through the PxIxIT docking site were found to play a central role in enhancing post-entry dengue virus infection.

Distinctions in Fc glycosylation among humans are an important driver of diversity in inflammatory responses. How basal Fc glycosylation impacts susceptibility to other infectious diseases, including during the neonatal period, is an essential topic for further discovery.

## Supplemental Figures & Tables

**Supplementary Figure 1.**
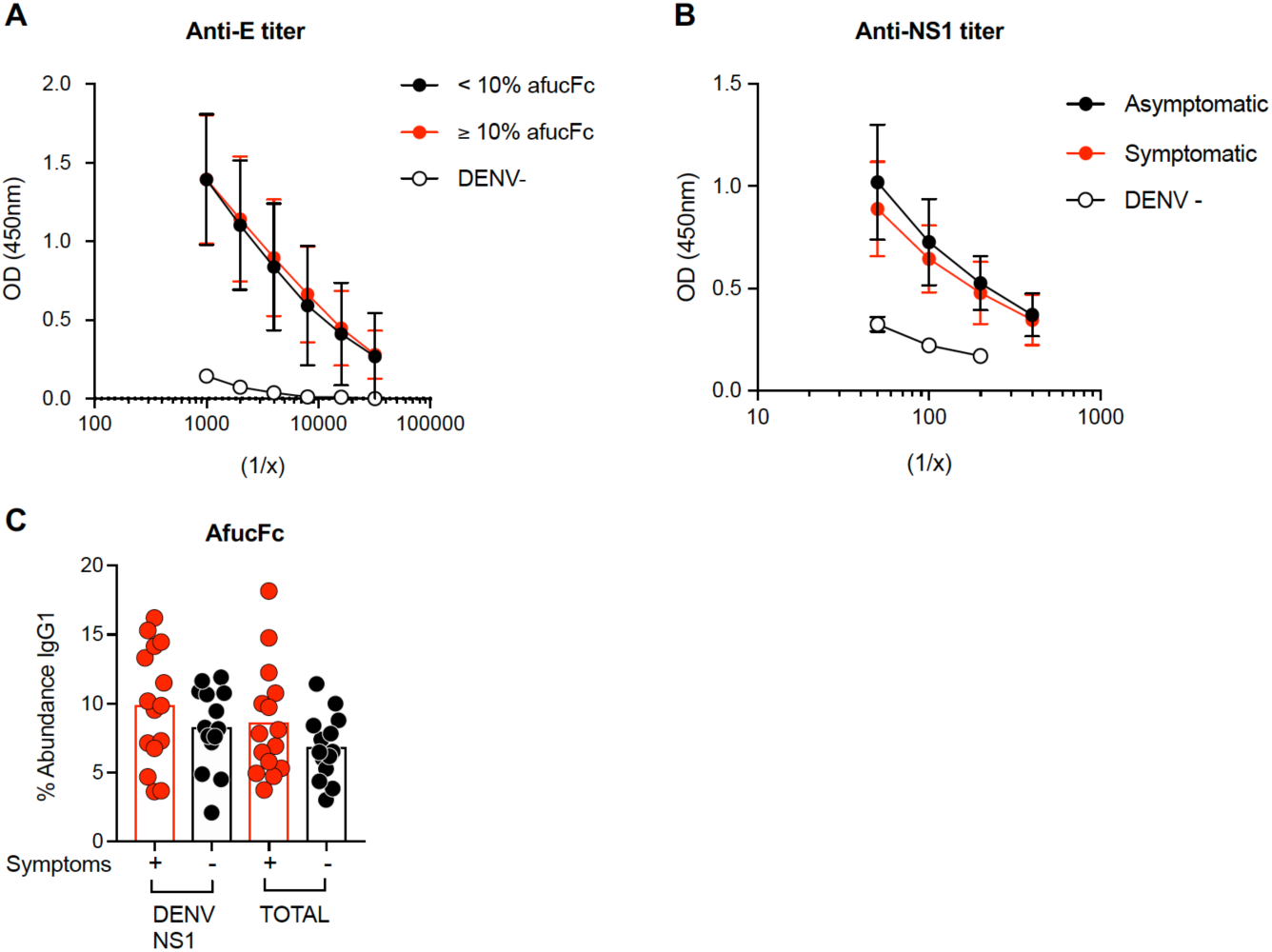
Characterization of anti-E IgGs in mothers of infants with symptomatic or asymptomatic primary dengue infections. **A)** The afucosylation status of anti-E IgGs did correlate with titer of IgGs that bound to dengue-infected cells **B)** Anti-NS1 IgG titer was not different between mothers of infants with symptomatic or asymptomatic primary dengue infections. **C)** Afucosylated glycoforms of NS1-reactive or total IgG were not distinct between mothers of infants with symptomatic or asymptomatic infections.

**Supplementary Figure 2.**
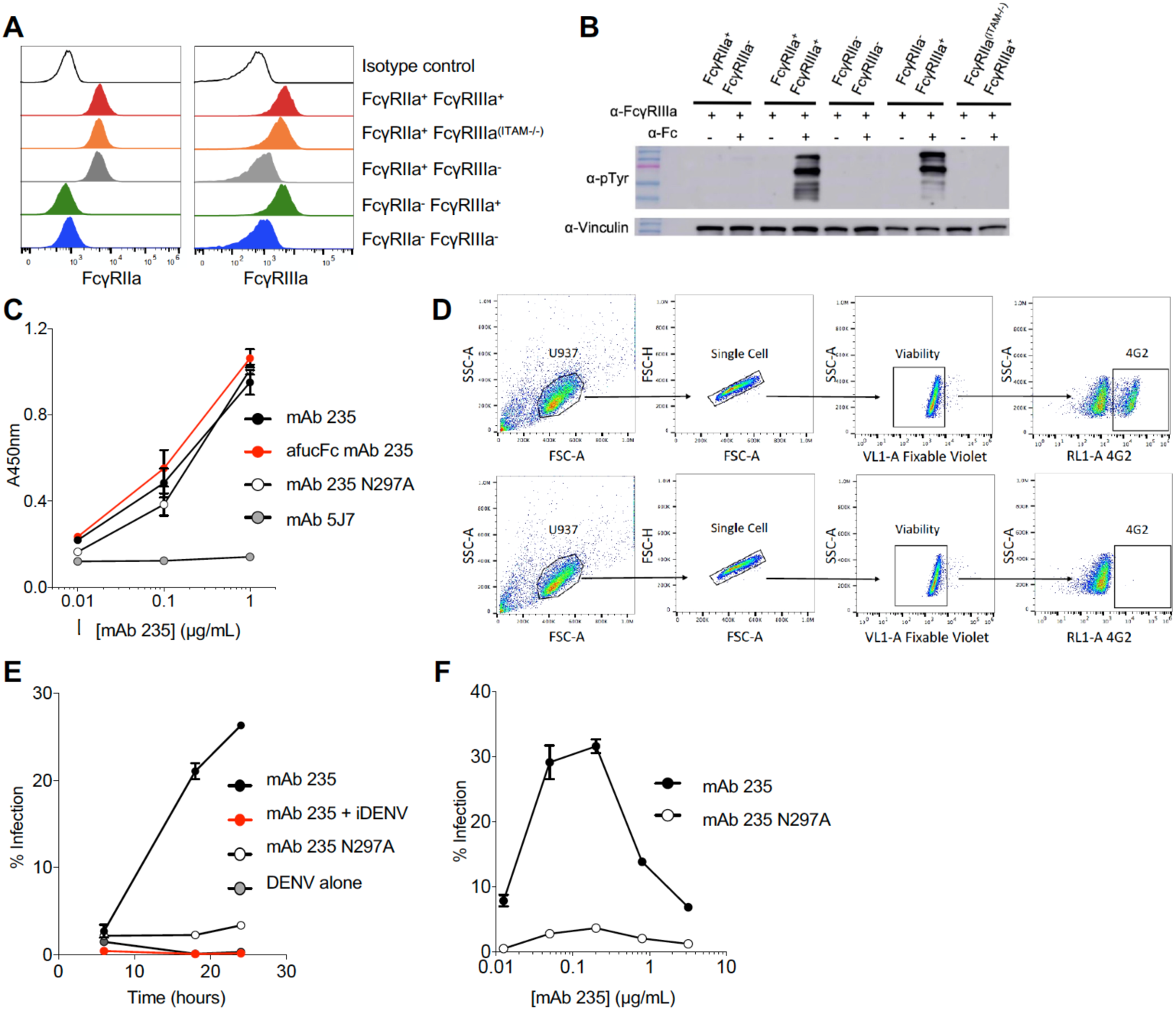
Engineered mAbs and human monocytes to study the role of Fc afucosylation and FcγRIIIa in antibody-dependent dengue infections. To define the roles of FcγRIIa and FcγRIIIa in infection by dengue immune complexes, a panel of human monocytic cell lines (U937) was generated that express all combinations of these activating FcyRs. **A)** Expression of FcγRIIa and/or FcγRIIIa on U937 cell lines was confirmed by FACS. **B)** Cross-linking of FcγRIIIa induced signaling in FcγRIIIa^+^ cells, indicated by tyrosine phosphorylation Signaling was observed in the FcγRIIa^+^FcγRIIIa^+^ and FcyRII^-^aFcγRIIIa^+^ cell lines but not the FcγRIIIa-deficient FcγRIIa^+^FcYRIIIa^(ITAM-)^ line, FcγRIIa^+^FcγRIIIa^-^(wild-type) or FcγRIIa^-^FcγRIIIa^-^cells. **C)** Broadly reactive anti-E mAb 235 was expressed as a human IgG1 (mAb 235), afucosylated Fc (afucFc) mAb235, mAb 235 Fc null variant N297A. The DENV3 specific mAb 5J7 was also expressed as a human IgG1. Binding of mAbs was assessed by ELISA on dengue 2-infected cells. **D)** FACS gating strategy for dengue-infected cells. Infection in single, live cells was assessed by staining of envelope protein using mAb 4G2 at 24 hours unless otherwise specified. **E)** DENV2 infection in wild-type U937 cells increased over time when infection occurred in the presence of mAb 235 expressed as a wild-type human IgG1. Infection did not increase when mAb 235 was expressed as an Fc-null variant, N297A or when immune complexes were made with deactivated virus (iDENV). **F)** The degree of enhanced infectability depended on the concentration of mAb 235.

**Supplementary Figure 3.**
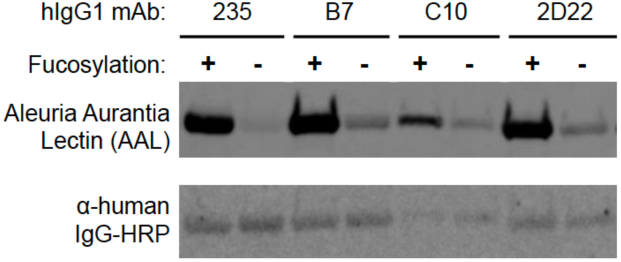
Blot showing afucosylation of dengue anti-E mAbs. Afucosylation of hIgG1 mAb 235, B7, C8, and 2D22 was confirmed by western blot. Fucosylation was detected with a biotinylated *Aleuria aurantia* lectin (AAL), detected with streptavidin HRP and anti-human IgG Fc HRP was used as a loading control.

**Supplementary Figure 4.**
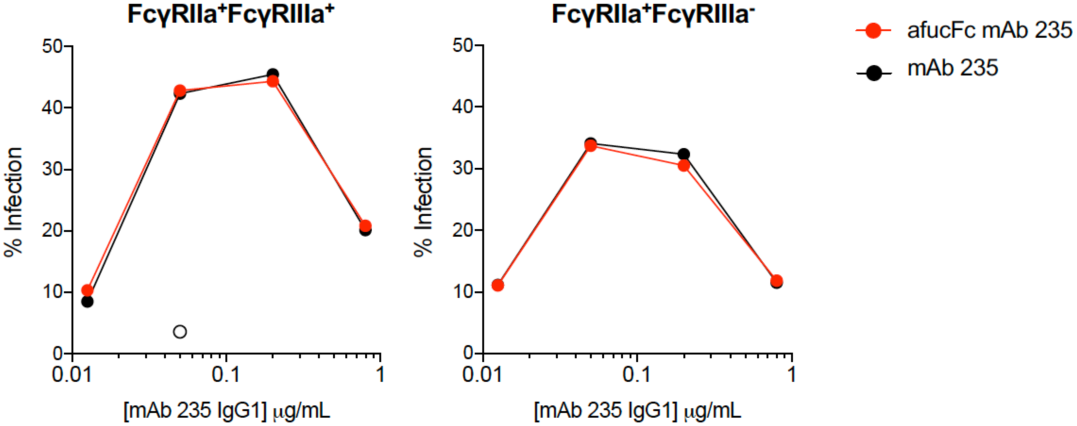
Afucosylation of anti-dengue IgGs does not impact infection in pure populations of FcγRIIa^+^FcγRIIIa^+^ or FcγRIIa^+^FcγRIIIa^-^ cells. Cells were infected with wild-type (fucosylated) mAb 235 IgG1 or afucosylated (afucFc) mAb 235.

**Supplementary Figure 5.**
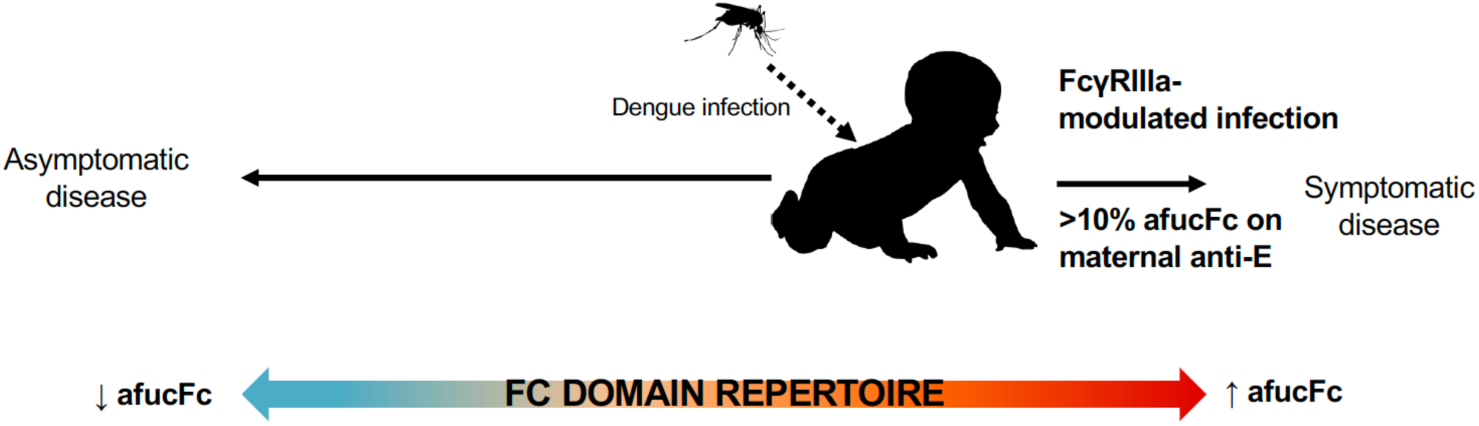
Model for susceptibility to clinically significant dengue disease. Primary dengue infection in infants of mothers with ≥10% in abundance of afucosylated anti-dengue Fc glycans is a risk factor for symptomatic infection due to modulation of FcYRIIIa-expressing cells during infection.

**Supplementary Table 1.** The binding epitopes and dengue-reactivity of anti-E mAbs 235, B7, 2D22, C10, C8, and 5J7. EDE: Envelope dimer epitope

| mAb Name | Alias | Epitope | Domain | Reactivity | Ref |
| --- | --- | --- | --- | --- | --- |
| 23-5G2D2 | 235 | E protein | DII | DENV2 | (Sasaki et al., 2013) |
| 747(4) B7 | B7 | Dimer | EDE2 | DENV1-4 | (Dejnirattisai et al., 2015; Rouvinski et al., 2015; Swanstrom et al., 2016) |
| 2D22 | 2D22 | Dimer | EDE3 | DENV2 | (de Alwis et al., 2012; Swanstrom et al., 2016) |
| 753(3) C10 | C10 | Dimer | EDE1 | DENV1-4 | (Rouvinski et al., 2015; Swanstrom et al., 2016) |
| 752-2 C8 | C8 | Dimer | EDE1 | DENV1-4 | (de Alwis et al., 2012; Rouvinski et al., 2015; Swanstrom et al., 2016) |
| 5J7 | 5J7 | E protein herring-bone | Quarternary epitope | DENV3 | (de Alwis et al., 2012; Fibriansah et al., 2015b; Swanstrom et al., 2016) |

## Materials and Methods

### Clinical Cohorts and Samples

Paired maternal and cord blood samples were obtained from women and infants enrolled in PROMOTE (NCT 02163447), a randomized clinical trial of novel antimalarial chemoprevention regimens in Eastern Uganda (Jagannathan et al., 2018). The study was approved by the Institutional Review Boards of the Makerere University School of Biomedical Sciences, the Uganda National Council for Science and Technology, and the University of California San Francisco. Written informed consent was obtained from all study participants. Samples from mothers of infants who had primary dengue infections were from a study that was approved by the institutional review boards of the Research Institute for Tropical Medicine, Philippines, and the University of Massachusetts Medical School. Mothers and their healthy infants were recruited and enrolled after providing written informed consent (ClinicalTrials.gov NCT00377754) (Libraty et al., 2009). Samples from the controlled influenza A virus challenge study have been previously described (Memoli et al., 2015). The study (ClinicalTrials.gov NCT01646138) was approved by the National Institute of Allergy and Infectious Diseases institutional review board and underwent an ethics review by the NIH Department of Bioethics prior to being conducted. The study was conducted in accordance with the provisions of the Declaration of Helsinki and Good Clinical Practice guidelines.

### Fc Glycan and IgG Subclass Analysis

Methods for relative quantification of Fc Glycans and IgG subclasses have been previously described (Wang et al., 2015; Wang et al., 2017). Briefly, IgGs were isolated from serum by protein G purification. Antigen-specific IgGs were isolated on NHS agarose resin (ThermoFisher; 26196) coupled to the protein of interest. Mixed envelope proteins from the four DENV serotypes were used (ProSpec; DEN-021, DEN-022, DEN-023, DEN-024). Following tryptic digestion of purified IgG bound to antigen-coated beads, nanoLC-MS/MS analysis for characterization of glycosylation sites was performed on an UltiMate3000 nanoLC (Dionex) coupled with a hybrid triple quadrupole linear ion trap mass spectrometer, the 4000 Q Trap (SCIEX). MS data acquisition was performed using Analyst 1.6.1 software (SCIEX) for precursor ion scan triggered information dependent acquisition (IDA) analysis for initial discovery-based identification.

For quantitative analysis of the glycoforms at the N297 site across the three IgG subclasses (IgG1, IgG2 and IgG3/G4), multiple-reaction monitoring (MRM) analysis for selected target glycopeptides, was applied using the nanoLC-4000 Q Trap platform to the samples which had been digested with trypsin. The m/z of 4-charged ions for all different glycoforms as Q1 and the fragment ion at m/z 366.1 as Q3 for each of transition pairs were used for MRM assays. A native IgGs tryptic peptide (131-GTLVTVSSASTK-142) with transition pair of, 575.9^+2^/780.4 was used as a reference peptide for normalization. IgG subclass distribution was quantitatively determined by nanoLC-MRM analysis of tryptic peptides following removal of glycans from purified IgGs with PNGase F. Here the m/z value of fragment ions for monitoring transition pairs was always larger than that of their precursor ions to enhance the selectivity for unmodified targeted peptides and the reference peptide. All raw MRM data was processed using MultiQuant 2.1.1 (SCIEX). All MRM peak areas were automatically integrated and inspected manually. In the case where the automatic peak integration by MultiQuant failed, manual integration was performed using the MultiQuant software.

### Generating U937 Cell Lines

Wild type U937 cells are FcγRIIa^+^FcγRIIIa^-^; FcγRIIa was deleted from the wild type line using CRISPR/Cas9 methods to generate FcγRIIa^-^ FcγRIIIa^-^ monocytes. Wild type or FcγRIIa^-^FcγRIIIa^-^ cells were stably transfected for expression of FcγRIIIa, thus generating the FcγRIIa^+^FcγRIIIa^+^ and FcγRIIa^-^FcγRIIIa^+^ lines, respectively. A final cell line, FcγRIIa^+^FcγRIIIa^(ITAM-/-)^, expresses FcγRIIa and a variant of FcγRIIIa that does not associate with the Fc receptor gamma chain, and thus lacks ITAM signaling (Blazquez-Moreno et al., 2017).

The gRNAs to target h FcγRIIa were designed utilizing DESKGEN software, and inserted into sequence containing U6 promoter and gRNA scaffold (Yang et al., 2014). The gRNA sequence to target human FcγRIIa was 5’GATGTATGTCCCAGAAACCTG 3’. The final fragment (Integrated DNA Technologies) was cloned into the pCR-BluntII-Topo vector. Human codon optimized cas9 DNA was obtained from (Addgene, Plasmid #41815). To generate the knockout cell lines, U937 cells were transfected with 1.5µg hCas9 and 0.5µg of gRNA via electroporation. The U937 cells were electroporated using Lonza Nucleofector 2b (program W-001) and the Nucleofection Kit C (Lonza, VPA-1004). After electroporation, U937 cells incubated at 37°C for 48 hours. Following incubation, cells were synchronized at the early S phase using a double thymidine block. The cells were stained with IV.3-FITC, and FcγRIIa negative cells were bulk sorted on SH800 sorter. After one bulk sort, the cells were single cell sorted based on negative FcγRIIa expression. The single cell clones were confirmed using Sanger sequencing. To generate FcγRIIIa^+^ cells, the wildtype cells were nucleofected with pLJM1-EGFP plasmid (Addgene, Plasmid #19319) that has EGFP replaced with FcγRIIIa or FcγRIIIa^(ITAM-/-)^, while the FcγRIIa^-^ cells were nucleofected with pLJMI-FcγRIIIa only. Stably transfected clones were selected using 1ug/mL puromycin.

### FACS staining of U937 cell lines

U937 cell lines were stained with a fixable live/dead stain (ThermoFisher; L10119). To assess FcγR expression cells were stained at 1:100 with a FITC conjugated anti-human FcγRIIa antibody, clone IV.3 (STEMCELL Technologies; 60012FI); Brilliant Violet 711 conjugated anti-human FcγRIII, clone 3G8 (BioLegend; 302044); an FcγRIIb antibody clone 2B6 (gifted by the Ravetch lab) conjugated to Alexa Fluor 647 (ThermoFisher; A20186) and an APC conjugated anti-human FcγRI, clone 10.1 (Biolegend, 305014) for 30 minutes at 4°C. The expression levels of CD32 and CD16 were measured using flow cytometry.

### Cross-linking of FcγRs

U937 cell lines were treated with mouse anti-FcγRIIIa/b, clone 3G8 (Biolegend; 302057) at 10 µg ml^−1^ in Optimem Medium on ice for 30 min. Anti-Mouse Fc was then added at 1.25 µg per 1 million cells and incubated at 37°C for 3 min to crosslink mab 3G8. Cells were then lysed in a one to one ratio of cell lysate buffer containing 100mM TRIS-HCL/300mM NaCL/2% Triton X 100 and a protease/phosphatase inhibitor cocktail (ThemoFisher; 88668). Protein concentrations were quantified using BCA Protein Assay Kit (ThermoFisher; 23227) and lysates were reduced by boiling for 5 min with Laemmli SDS sample buffer containing 5% β-mercaptoethanol. The lysates were run on Mini-PROTEAN® TGX™ Precast Protein Gels (Bio Rad; 456903). Semidry transfer to PVDF membranes was performed. Blots were blocked 5% BSA in 1x TBS buffer 20mM Tris-HCL/150mM NaCl pH 7.6 for 2 hours and were subsequently incubated with anti-phosphotyrosine secondary antibody (Biolegend; 309302) at 1:1,000 in 5% BSA in TBS, 0.2% Tween-20 (TBST) overnight at 4 ° C. Blots were washed and bands were detected with anti-mouse IgG Fc-HRP (Southern Biotech; 1030-05) at 1:10,000 for 1 hour at room temperature. Bands were detected with chemiluminescent substrate Western Lightning ECL Pro (Perkin Elmer; NEL120001EA) according to manufacturer’s instructions. The membrane was then imaged using the Amersham Imager 600 (GE Life Sciences) with auto exposure under chemiluminescent detection settings. Post-imaging, the membrane was washed extensively with 1x TBST before treating with Restore™ Western Blot Stripping Buffer (ThermoFisher; 21059) for 15 min at room temperature. To analyze protein loading, the membrane was re-blocked with 5% BSA in 1x TBST for 2 hour at room temperature before probing with a 1:200 dilution anti-vinculin 7F9 (Santa Cruz Biotechnology; sc-73614) in 5% BSA/1x TBST for 1 hour at room temperature followed by anti-mouse IgG HRP secondary, as above. The membrane was then treated with chemiluminescent substrate, and imaged as described above.

### Antibody Expression

Antibodies: 23-5G2D2747(4)(Sasaki et al., 2013), B7 (B7)(Dejnirattisai et al., 2015; Rouvinski et al., 2015; Swanstrom et al., 2016), 2D22(de Alwis et al., 2012; Fibriansah et al., 2015a; Swanstrom et al., 2016), and 753(3) C10 (C10)(Rouvinski et al., 2015; Swanstrom et al., 2016), 752-2 C8(de Alwis et al., 2012; Rouvinski et al., 2015; Swanstrom et al., 2016), and 5J7(de Alwis et al., 2012; Fibriansah et al., 2015b; Swanstrom et al., 2016) were expressed in Expi293 cells as follows. For afucoyslated variants, Expi293 cells (ThermoFisher; A14527) were split to 2.0 x 10^6^ cells/mL in a 10 mL volume 24h prior to transfection and treated with 100 μM of a fucosyl transferase inhibitor, 2F-Peracetyl-Fucose (Rillahan et al., 2012) (EMD Millipore, 344827). On the day of transfection, 5μg filter-sterilized DNA (2.5μg each of heavy and light chain-encoding plasmids) was added to 1mL of Expi293 medium (ThermoFisher Scientific; A1435101) in a fresh tube, and mixed. To this, 13μL (1.3μL/mL of total transfection volume) of FectPro transfection Reagent (Polyplus Transfection; 116-001) was added. The DNA-transfection reagent mixture was incubated at room temperature for 10 minutes before adding dropwise to 10 mL of the Expi293 cells. The cells were then additionally treated with a final concentration of 4g/L glucose and 3 mM valproic acid. The cells were then grown under shaking conditions (125 rpm) at 37°C for 3 days before receiving an additional 4g/L final concentration of glucose. The cells were harvested on day 6 post-transfection, the cell pellet discarded, and the supernatant media filtered before purification. MAbs were purified over Protein G Sepharose 4 Fast Flow resin (GE Healthcare; 17061802). The antibodies were buffer exchanged into PBS pH 7.4 prior to use in any cell based assays.

### Cell-based ELISA

Wells of a 96 well plate were seeded with 3×10^4^ Vero cells (ATCC; CCL-81). The next day cells were infected with DENV2 New Guinea C (BEI Resources; NR-84) or DENV3 C0360/94 (BEI Resources; NR-48800) MOI 1 for 1 hour, washed, and incubated for 48 h in MEM, 1% methylcellulose, 2% inactivated FBS, antibiotics, and L-glutamine. Cells were fixed with 4% formaldehyde for 30 minutes and permeabilized in 2% inactivated FBS/0.1% saponin/1xPBS. For analysis of clinical samples, serially diluted maternal sera were incubated for 2 hours at room temperature on cells infected with the virus serotype of the paired infant’s dengue virus infection. For mAb binding assays, mAbs were titrated and incubated for 2 hours at room temperature on DENV2 infected cells. Cells were washed with PBS–0.1% Tween 20 and incubated with mouse anti-human IgG Fc-HRP (Southern Biotech; 9040-05) for 1 hour. Following incubation, the wells were washed and TMB Liquid Substrate (Sigma; T0440) was added and stopped with 1M HCL after approximately 5 minutes. Absorbance at 450 nm was measured; ELISA performed in triplicate.

### NS1 ELISA

Wells of a 96 well plate were coated with dengue anti-NS1 protein (ProSpec DEN-004) for 1h at 37°C and blocked in 1% BSA/PBS for 1h at 37°C. For analysis of clinical samples, serially diluted maternal sera in 1% BSA/PBS were incubated for 1 hours at 37°C on NS1 coated plates. Plates were washed with PBS–0.1% Tween 20 and incubated with mouse anti-human IgG Fc-HRP (Southern Biotech; 9040-05) for 1 hour. Following incubation, the wells were washed and TMB Liquid Substrate (Sigma; T0440) was added and stopped with 1M HCL after approximately 4 minutes. Absorbance at 450 nm was measured; ELISA performed in triplicate.

### Antibody Dependent U937 Infections

Anti-E mAbs were incubated with dengue virses for 30 minutes at 37°C to form immune complexes. 5×10^4^ U937 cells were added to the immune complexes (MOI: 4) in RPMI containing 2% inactivated FBS, antibiotics, MEM nonessential amino acid solution, 2-mercaptoethanol, and HEPES. Cells were infected for 2 hours at 37°C, washed, then resuspended in infection media and incubated for an additional 4-24 h at 37°C. Infected cells were stained with a fixable violet live/dead stain (ThermoFisher; L34955) for 30 min, fixed with 4% formaldehyde and permeabilized in 2% inactivated FBS, 5mM EDTA, 0.1% saponin in 1xPBS. Cells were stained using an anti-E antibody (4G2; ATCC) conjugated to Alexa Fluor 647 (ThermoFisher; A20186) at 1:400 for 30min. The frequency of infected cells was determined using flow cytometry, defined as the percentage of live, single cells that were positive for mAb 4G2 staining. For infections involving inhibitors of ITAM signaling, U937s were treated with titrated inhibitors: PRT062607 (Fisher Scientific; NC0664362), INCA-6 (Abcam; ab145864), 11R-VIVIT (Thermo Fisher; CM102281.1), or 11R-VEET (Thermo Fisher; CM102281.2) in complete RPMI media for 2h at 37C prior to, during or following infection as indicated. Immune complexes were formed at 37°C for 30min between hIgG1 mAb 235 and DV2 (MOI4) at 37°C. U937 were washed 2 times with PBS and then added to the immune complexes for a 2h infection at 37°C. Cells were then washed 2x in PBS and re-suspended in infection media followed by incubation for an additional 22 h at 37°C. For inhibition of PMA-ionomycin treatment: U937s were treated with titrated inhibitors 11R-VIVIT or 11R-VEET in complete RPMI media for 2h at 37C. Cells were then washed 2 times followed by treatment with PMA (Invivogen; tlrl-pma) and 50ug/ml ionomycin (Invivogen; inh-ion) in complete RPMI media for 6 h at 37°C prior to infection with dengue immune complexes.

### Blocking Assay

Anti-E monoclonal antibody 235 expressed as a human IgG1 (hIgG1) was incubated with dengue virus for 30 minutes at 37°C to form immune complexes and then cooled on ice for 15 min. FcγRIIa^+^FcγRIIIa^+^ U937 cells were incubated on ice for 30 minutes with different combinations of FcγR-specific blocking antibodies, each at 4ug/ml. Blocking antibodies: FcγRIIa antibody, clone IV.3 (Stem Cell Technology Catalog; 60012); anti-human FcγRIIIa, clone 3G8 (Biolegend; 302057); and anti-human FcγRI, clone 10.1 (Biolegend, 305016). Total antibody concentration between determinants was controlled by addition of human IgG2/kappa where needed (Sigma Aldrich; AG504). Cells were washed with ice-cold PBS and added to dengue immune complexes (MOI 5). Infection proceeded for 1 hour on ice. Cells were then washed and resuspended in infection media followed by incubation for an additional 22 h at 37°C.

### Avidity ELISA

The avidity ELISA has been described before (Wang et al., 2015). Briefly, Wells of microtiter plates were coated with DENV2 (ProSpec; DEN-022) or DENV3 (ProSpec; DEN-023) E proteins (2.5µg ml^-1^ in PBS) for 2 hours at 37°C. Plates were blocked for 30 min with 1% BSA/1%PBS and then incubated with titrated Gentle Ag/Ab Elution Buffer, pH 6.6 (ThermoFisher; 21027) and maternal IgG, which was normalized for optical density of binding to dengue-infected cells and matched for virus serotype of the paired infant’s dengue virus infection. Plates were washed with PBS/0.1% Tween 20 and wells were incubated with alkaline phosphatase-conjugated anti-human Fc (Southern Biotech; P7998) followed by 2 hour incubation. Following incubation, the wells were washed and p-nitrophenyl phosphate substrate solution (Sigma; P7998) was solution was added. Absorbance at 405 nm was measured after approximately 45 min; ELISA performed in triplicate.

### Western Blot of Antibodies

Antibodies were run under reducing and denaturing conditions by SDS-PAGE on a gradient gel with Tris-Glycine buffer. The gel was transferred to a 0.45μm nitrocellulose membrane (Bio-Rad) using semi-dry transfer. The membrane was blocked in 5% BSA in 1x TBST for 2 hours at room temperature, before probing with a 1:10,000 dilution of biotinylated *Aleuria Aurantia* Lectin or AAL (Vector Labs; B-1395) in 5% BSA/1x TBST for 1 hour at room temperature. After 4-6 washes with 1x TBST, the membrane was incubated with a 1: 2500 dilution of streptavidin-HRP (Southern Biotech; 7100-05) in 5% BSA/1x TBST for 1 hour at room temperature. After 4-6 washes with 1x TBST the membrane was treated with chemiluminescent substrate Western Lightning ECL Pro (Perkin Elmer; NEL120001EA) according to manufacturer’s instructions. The membrane was then imaged using the Amersham Imager 600 (GE Life Sciences) with auto exposure under chemiluminescent detection settings. Post-imaging, the membrane was washed extensively with 1x TBST before treating with Restore™ Western Blot Stripping Buffer (ThermoFisher; 21059) for 15 min at room temperature. To analyze protein loading, the membrane was re-blocked with 5% BSA in 1x TBST for 2 hours at room temperature before probing with 1:5000 dilution of anti-human IgG-HRP (Southern Biotech) in 5% BSA/1x TBST for 1 hour at room temperature. The membrane was then treated with chemiluminescent substrate, and imaged as described above.

## Acknowledgments

We thank Martha S. Cyert (Stanford University) for helpful discussions. The authors also thank the Infectious Diseases Research Collaboration and the PROMOTE study team (NICHD PO1HD059454, including Drs. Diane Havlir, Margaret Feeney, and Grant Dorsey (UCSF), and Prof Moses Kamya (Makerere University) for provision of paired maternal and cord blood clinical samples. Support was received from Stanford University, the Chan Zuckerberg Biohub and the Searle Scholars Program. Research reported in this publication was supported in part by the Bill & Melinda Gates Foundation (OPP1188461) and the National Institutes of Health (5K22AI12347802, 5U19AI111825-05). The influenza A virus challenge study was supported by the Intramural Research Program of the National Institute of Allergy and Infectious Diseases (NIAID).

